# Prioritization over certainty: Action planning in visual working memory is coupled with selective attention, rather than action predictability

**DOI:** 10.64898/2026.11.18.767539

**Authors:** Şahcan Özdemir, Nursena Ataseven, Wouter Kruijne, Elkan G. Akyürek, Daniel Schneider

**Affiliations:** Leibniz Research Centre for Working Environment and Human Factors, Dortmund, Germany; Department of Experimental Psychology, University of Groningen, Groningen, the Netherlands

**Author notes:** Corresponding Author Şahcan Özdemir.

**Keywords:** working memory, action planning, selective attention

## Abstract

When visual information is no longer perceptually available, working memory (WM) allows us to act on sensory representations. But does this guidance require attentional prioritization? We addressed this question by asking participants to remember two oriented bars that were later tested with one predictable and one random probe. This design allowed us to track when a remembered orientation became linked to a specific response hand and whether that link depended on which item was currently prioritized in WM. Response preparation followed prioritization: even when the later response was predictable, no motor preparation emerged while the corresponding item remained latent. Our exploratory analyses showed that prospective action information reduced the need to reprioritize latent items, and that learned stimulus–response mappings generalized across task contexts, suggesting that task-relevant features of visual WM can be partly offloaded onto motor codes. These findings show that active response preparation is coupled with selective attention, while motor representations can also support maintenance in a visual WM task.

## Introduction

Complex behavior requires us to integrate and manipulate sensory information, plan possible actions, select among them, and execute the chosen response. Consider a basketball player who receives the ball and must rapidly decide to pass the ball to one of the teammates. The player may have to rely on visual information that is temporarily unavailable in perception, such as the location of a teammate who has moved out of sight. When relevant sensory information is no longer accessible, working memory (WM) provides a fundamental cognitive mechanism for briefly maintaining that information to support goal-directed behavior (Baddeley, 2003, 2010).

More specifically, the maintained task-relevant information can guide the planning and subsequent execution of overt actions (Boettcher et al., 2021; Rösner et al., 2022; Schneider et al., 2017; van Ede, 2020; van Ede et al., 2019). In the basketball example, the motor movements required to pass the ball can be represented and maintained as a potential action plan. However, the mechanisms governing the relationship between visual representations and action plans remain unclear. Does preparing a pass require selectively attending to the intended teammate?

In this study, we investigate whether the attentional mechanisms underlying visual prioritization and response preparation are dependent or can operate independently. If these processes are coupled, prioritizing associated action plan should require a prioritized item in visual WM. Alternatively, if visual and action-related representations can be prioritized independently, an action plan may be prepared and maintained even when attention is directed toward a different WM representation. Such flexibility may be possible when an upcoming response can be anticipated in advance, allowing action preparation to proceed independently of the current focus of selective attention.

Olivers and Roelfsema (2020) proposed that selective attention links WM representations to the overt actions or covert mental operations that are currently relevant for behavior. From this perspective, prioritizing a visual WM item should also prioritize the action associated with that item. Consistent with this account, previous studies have shown that response selection emerges together with the prioritization of task-relevant information (Echeverria-Altuna et al., 2025; Özdemir et al., 2025; Rösner et al., 2022; van Ede et al., 2019; Zickerick et al., 2021), according to the task demands (Heuer et al., 2020; Myers et al., 2017). These findings suggest that attentional selection may jointly prioritize the visual representation in WM and the associated action plan. Accordingly, this account predicts that response preparation is coupled to selective attention, such that selectively prioritizing a WM item for an upcoming task also prioritizes its associated response.

Alternatively, WM may support response preparation and selection more flexibly. Depending on the available information and task demands, WM may rely on different mechanisms, including maintaining information in different representational formats (Balaban & Luria, 2016; Yang et al., 2026), shifting between long-term memory and WM representations (Özdemir et al., 2024; Senturk et al., 2023), and using activity-silent mechanisms to maintain latent information (Rose et al., 2016; Stokes, 2015; Wolff et al., 2017). Extending this flexibility to action planning, the predictability of an upcoming action may be a critical factor determining whether response preparation depends on the prioritization of the corresponding visual WM item. Specifically, when the required action can already be anticipated from the task context, its preparation may proceed independently of attentional prioritization. Under these circumstances, an action plan may remain prioritized or undergo further preparation even when selective attention is directed toward another WM item. Thus, action predictability may allow WM to decouple motor preparation from the selective prioritization within visual WM.

We designed a WM task that allowed us to dissociate the attentional prioritization of WM items from the preparation of their associated responses by manipulating response predictability and test order. Participants memorized two tilted bars in WM and subsequently reported each orientation using a probe whose starting orientation was either predictable or randomly determined. For the predictable probe, each item’s tilt could be mapped in advance onto a left-or right-hand response. A retro-cue presented during the maintenance interval indicated which item would be tested first, while the order of the predictable and unpredictable probes was manipulated across blocks. Critically, when the predictable probe was scheduled for the second test, the response required for that later-tested item could already be anticipated while visual attention was directed toward the item tested first. This condition therefore allowed us to test whether an action plan could be prepared despite the corresponding visual WM item being uncued.

## Results

Thirty-four human subjects participated in the study. Thirty-two participants were included in the final EEG analysis, and thirty-three in behavioral analysis (for the details of the exclusion criteria see Materials and Methods). Participants performed a visual WM task (Figure 1), while EEG was recorded. In each trial, they remembered the orientations of two tilted bars in distinct colors (orange and blue). Subsequently, a retro-cue, which was a colored central dot, was presented in either orange or blue color, indicating that the bar matching its color would be tested first. The uncued bar would then be tested second. Testing was done by means of two successive probes that the participants had to rotate clockwise (CW) or counterclockwise (CCW), to reproduce the corresponding memory items. The rotation directions were mapped to corresponding hands (CW, right-hand; CCW, left-hand), allowing us to measure lateralized preparation indices for each item separately. Critically, the predictability of each probe’s initial orientation was manipulated across blocks; in half of the blocks, the starting orientation of the first probe was predictable (i.e., vertical orientation) and the second probe was presented with a random orientation. In the other half, the first probe was random and the second probe was predictable (for a more detailed explanation of the experimental procedure, please see Materials and Methods).

**Figure 1.**
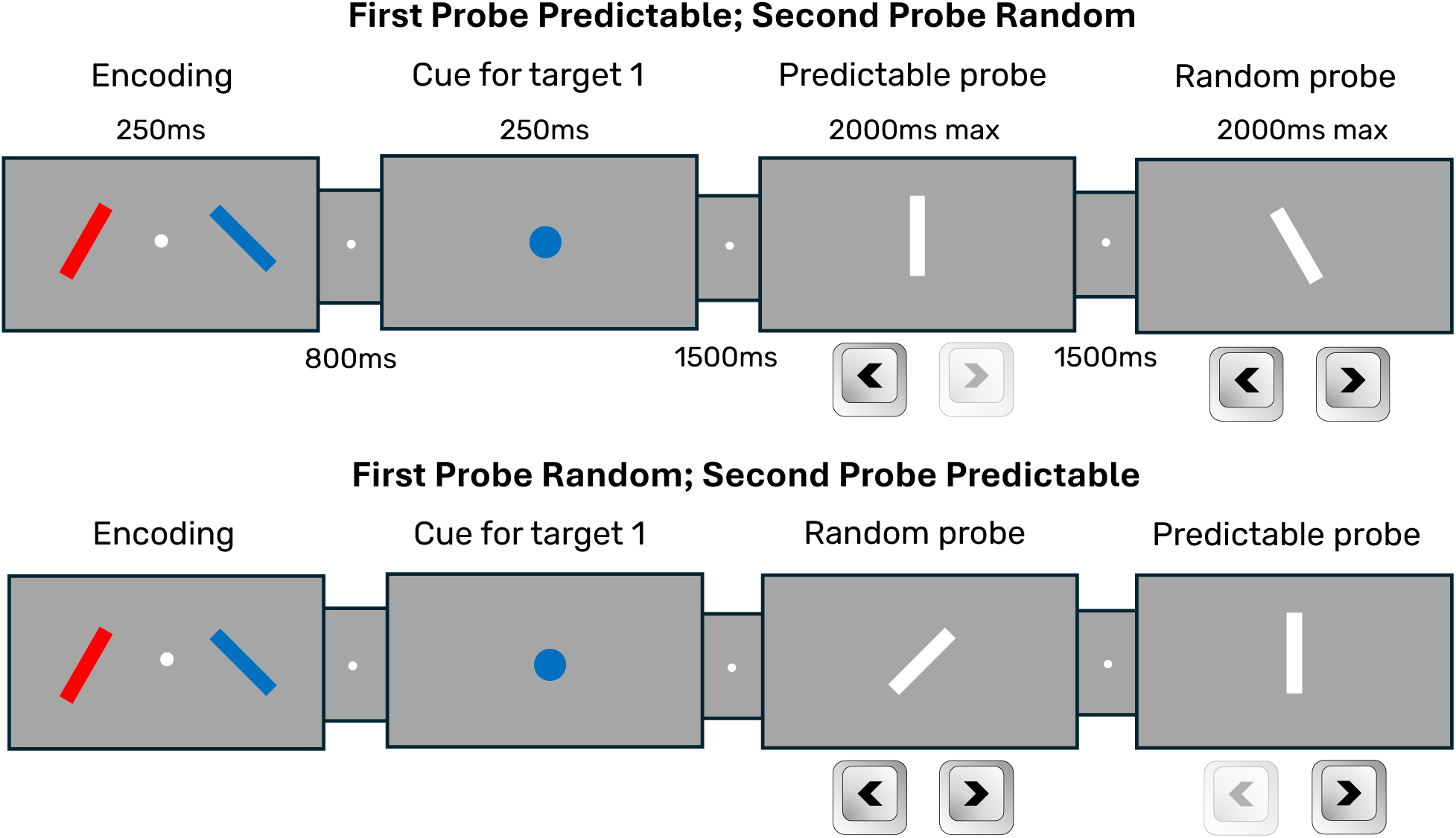
Experimental Design. The experiment consisted of a delayed match-to-sample task. On each trial, participants remembered two orientations. A color cue indicated which orientation would be tested first, while the other orientation was always tested second. One of the test probes always appeared in a vertical position (predictable probe), and participants were informed at the beginning of each block whether the predictable probe would appear first or second. For the predictable probe, participants were required to respond with the hand corresponding to the tilt direction of the target orientation. For the random probe, participants could use whichever response hand was most convenient given the probe position.

### Action plans benefit memory performance for the uncued item

First, we examined whether probe predictability benefited WM performance (for details of the performance measure, see Materials and Methods). As shown in Figure 2A, performance was better for the first than the second test (*F*(1,32) = 146.892; *p* < .001; *η2* = 0.473; *BF_incl_* = 3.061 x 10^10^) and for predictable than random probes (*F*(1,32) = 76.003; *p* < .001; *η2* = 0.231; *BF_incl_*= 7.904 x 10^6^). These effects interacted, with a larger benefit of probe predictability for the second test (*F*(1,32) = 38.182, *p* < .001, *η²* = 0.052, *BF_incl_* = 79.271 × 10³).

**Figure 2.**
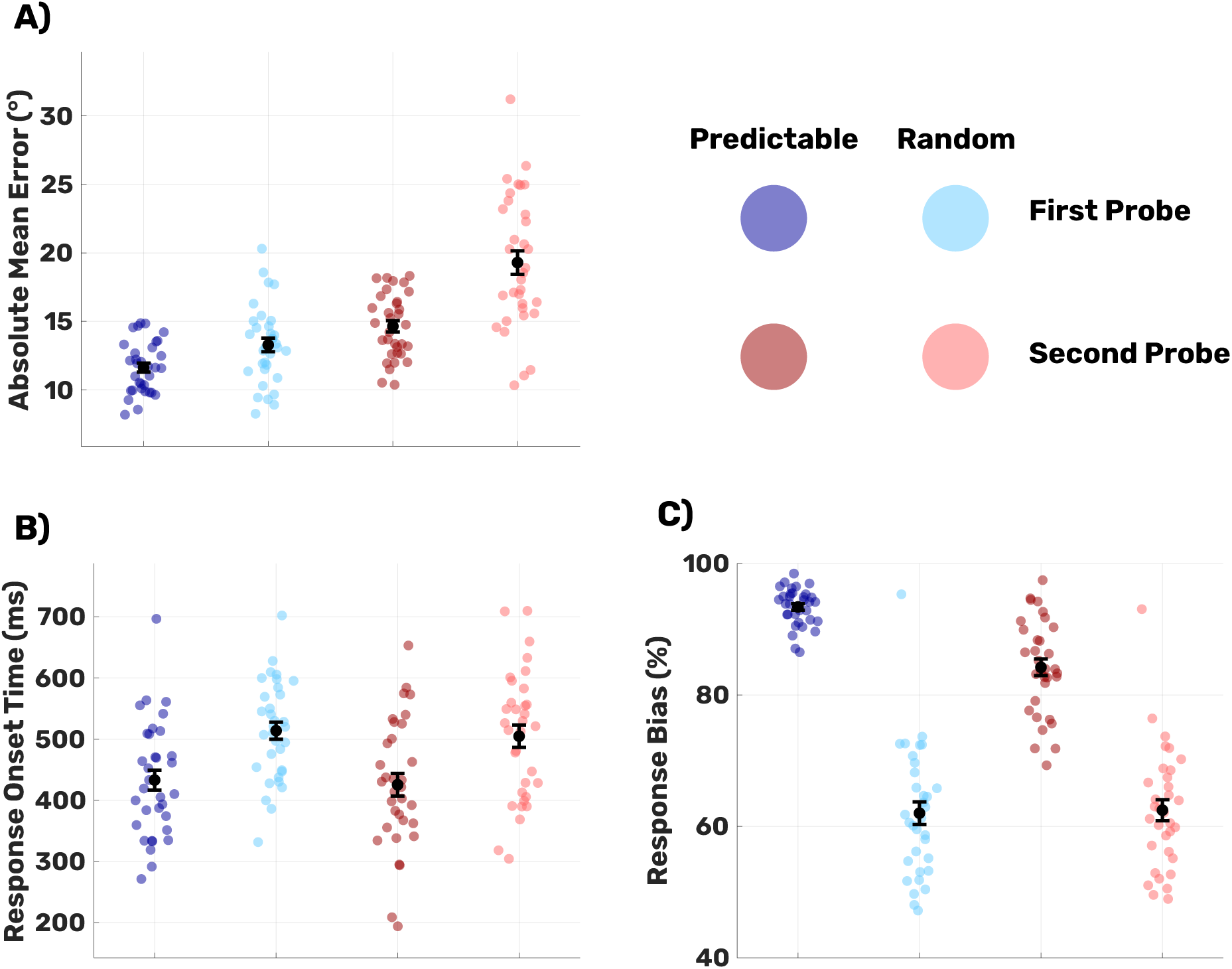
Behavioral comparisons. (A) shows comparisons of memory performance across all four conditions. Memory performance was quantified as the mean absolute error relative to the target orientation. (B) shows comparisons of response onset times across conditions. (C) shows comparisons of response bias. Response bias was calculated as the percentage of trials in which participants used the hand corresponding to the target orientation’s tilt direction in each condition.

Because the predictable probe always allowed the shortest rotation toward the target, we tested whether this response advantage accounted for the performance benefit. We therefore recalculated performance using only random-probe trials in which participants also applied the shortest rotation option. Performance remained better for the first than the second test (*F*(1,32) = 148.106; *p* < .001; *η2* = 0.687; *BF_incl_* = 7.437x10^10^). Critically, probe type interacted with test order, and the interaction (*F*(1,32) = 13.791, *p* < .001; *η2* = 0.021; *BF_1incl_*= 55.371) indicated that predictable probes improved performance only when they occurred second (*t*(32) = 2.999; *p_holm_* < .01; *d* = 0.333; 95% CI [0.079 1.677]), but not when they occurred first (*t*(32) = 1.418; *p_holm_* =.162; *d* = 0.157; 95% CI [0-0.384 1.214]). Accordingly, there was no overall main effect of probe type in this restricted analysis (*F*(1,32) = 0.966, *p* = .333; *η2* = 0.003; *BF_incl_*= 0.335). In line with the initial analysis, this pattern shows that the benefit of probe predictability was concentrated on the second-tested item, indicating that prospective action information is especially beneficial when the corresponding WM item is reprioritized from a latent state.

Probe predictability also reduced response onset times (Figure 2B), with faster responses for predictable than random probes (Figure 2B; *F*(1,32) = 41.631; *p* < .001; *η2* = 0.454; *BF_incl_* = 29.859 x 10^3^). Response onset times showed neither a main effect of test order (*F*(1,32) = 1.063; *p* = .310; *η2* = 0.0005; *BF_incl_* = 0.381) nor an interaction between probe type and test order (*F*(1,32) = 0.025; *p* = .876; *η2* = 3.482 x 10^-5^; *BF_incl_* = 0.324).

### Stimulus characteristics can lead to early action planning at encoding

Before moving onto the retro-cue window and within-WM prioritization, we first tested whether we observe early response preparation at encoding stage regarding the learned tilt-response hand matching (Nasrawi et al., 2023). We tested this in a subset of trials in which both items were tilted in the same direction. In these trials, both WM items afforded using the same tilt-matching hand for the predictable probe (right hand for clockwise items; left hand for counterclockwise items), regardless of which item was subsequently cued and tested first. Thus, the prospective response hand could already be selected during encoding based on stimulus tilt.

Suppression of mu/beta (8–30Hz) oscillatory power over centro-parietal electrodes contralateral to the response hand is a well-established neural marker of response preparation (Boettcher et al., 2021; Özdemir et al., 2025; Schneider et al., 2017; for calculation see Materials and Methods). We therefore treated the tilt-matching hand as the prospective response hand and compared contralateral with ipsilateral oscillatory power using a cluster-based permutation test (CBPT; for analysis details, see Materials and Methods).

When both items were tilted in the same direction, we observed mu/beta lateralization during encoding, between target onset and retro-cue presentation. A CBPT conducted from stimulus onset (0ms) to cue onset (1050ms) revealed response preparation both when the predictable probe was tested first (Figure 3A; between 244-1052ms and 7.6-17Hz; cluster size = 786, *d_max_* = -0.684; *d_avg_*= -0.45) and when the random probe was tested first (Figure 3B; between 384-1052ms and 7.6-27.6Hz; cluster size = 1067, *d_max_* = -0.828; *d_avg_*= -0.499). These findings showed that stimulus–response mapping can elicit response preparation already during encoding, even before the upcoming retro-cue provides probe-specific anticipatory information.

**Figure 3.**
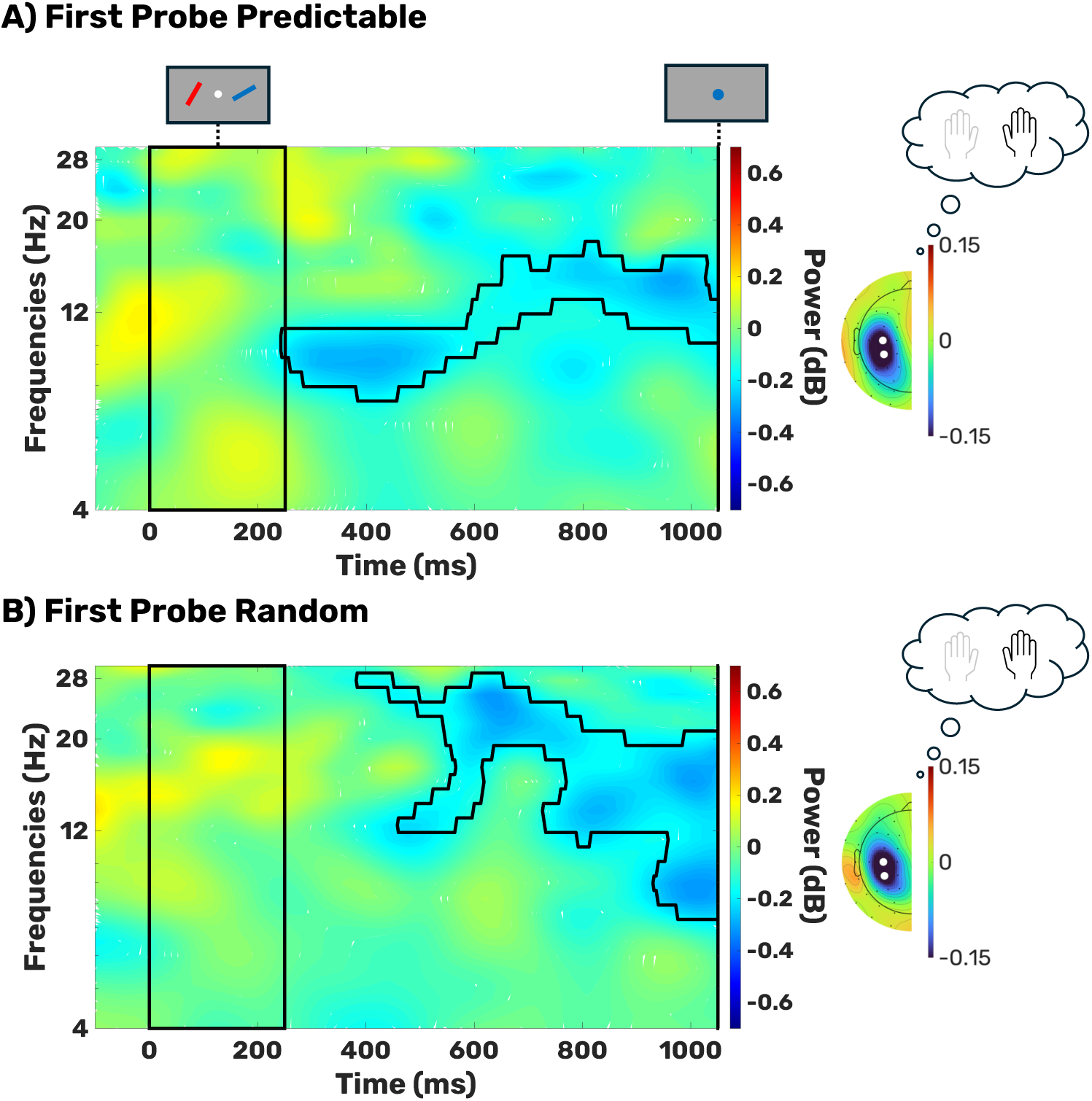
Response planning following WM encoding. The figure shows lateralization of the mu/beta (8-30Hz) power in EEG regarding the tilt-matching hand (channels: C3, C4, CP3 and CP4), for trials in which both items were tilted in the same direction. In all time-frequency figures, black outlined areas indicate significant clusters identified through CBPT. White dots on each topography represent the selected channels used for the comparison. For illustration purposes, the lateralization topoplots are depicted as lateralized against the right hemisphere; the calculation was always lateralized against the tilt-matching hand. Rectangular boxes represent the stimulus presentation window (onset at 0ms and offset at 250ms), both figures end with the cue presentation. Regardless of whether the first probe was predictable (A) or random (B), participants prepared the response hand that matched the items’ tilt at the encoding stage.

### Selective attention drives the transition from sensory WM representations to overt action preparation

If attentional prioritization is required to link a WM item to its upcoming response, we predicted that mu/beta lateralization associated with each item should emerge when that item becomes relevant to the upcoming probe. We therefore conducted a CBPT from cue onset (1050ms) through the post-response period (4100ms). As shown in Figure 4, response preparation emerged when the cued item became relevant for the upcoming test. This pattern was observed for predictable first probes (Figure 4A; 1396–4100ms, 4.3–30Hz; cluster size = 8934, d_max_ = −1.256, d_avg_ = −0.632) and also for random first probes (Figure 4B; 1300–4100ms, 7.6–30Hz; cluster size = 7238, d_max_ = −1.264, d_avg_ = −0.654), with mu/beta lateralization corresponding to the tilt-matching hand emerging before probe presentation. The two conditions differed only after probe onset (Figure 4C; 3116–4100ms, 4–14.5Hz; cluster size = 2226, d_max_ = −0.844, d_avg_ = −0.037). This post-probe divergence is consistent with the fact that the random probe could require a response hand different from the tilt-matching hand, whereas the predictable probe preserved the anticipated mapping. Together, these results supports the notion that response preparation follows attentional prioritization of the relevant WM item.

**Figure 4.**
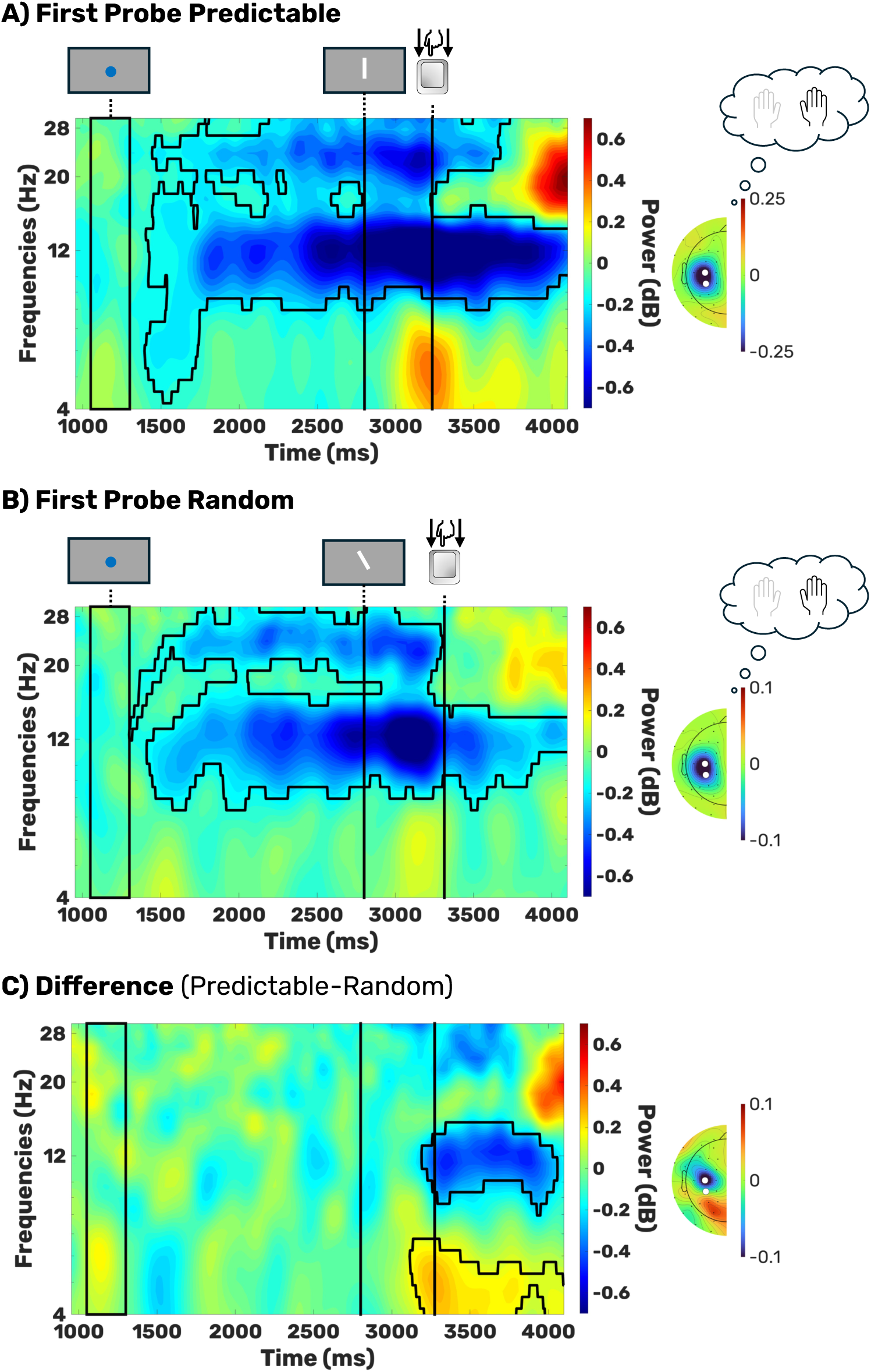
First response planning following retro-cue. The figure shows lateralization of mu/beta (8–30Hz) EEG power with respect to the tilt-matching hand (channels: C3, C4, CP3, and CP4). The rectangular box indicates the cue presentation window (1050–1300ms). The first black vertical line indicates probe onset, and the second black vertical line indicates the average response onset (button press). In A and B, the response line represents the average response onset within each condition, whereas in C it represents the average response onset across all trials. Regardless of whether the probe was predictable (A) or random (B), significant clusters of EEG power lateralization emerged with respect to the prepared response hand. C shows a difference in lateralization between the predictable and random probe conditions following the response.

### Response preparation does not occur independently of attentional prioritization, even with action predictability for the uncued item

When the second test used the predictable probe, participants could in principle prepare the response for the second-tested item after the retro-cue and before the first memory test. Because the first-tested item was paired with a random probe, no prospective response information was available for the currently prioritized item. This condition therefore allowed us to test whether response preparation for the latent second-tested item could occur while selective attention was directed toward the first item. We calculated mu/beta lateralization relative to the second-tested item’s tilt-matching hand and tested it from retro-cue onset to the onset of the first probe. The CBPT revealed no significant mu/beta lateralization over centro-parietal electrodes during this interval (1050–4100ms). Thus, even when the upcoming response for the second-tested item was predictable, we found no evidence that its action plan was prepared while the item remained outside the focus of attention.

We next examined when response preparation for the second probe emerged after completion of the first response. This analysis was time-locked to the onset of the second probe, which occurred 1500ms after the first response and therefore varied across trials. A CBPT conducted from −1500ms to 1500ms relative to second-probe onset showed clear response preparation for the second-tested item, regardless of whether the second probe was predictable (Figure 5A; −1500–1500ms, 7.6–30Hz; cluster size = 7867, dmax = −1.081, davg = −0.611) or random (Figure 5B; −1500–1500ms, 4–30Hz; cluster size = 9399, dmax = −1.230, davg = −0.600). Together with the absence of response preparation for the second-tested item during the first maintenance period, these findings suggest that even when its upcoming response could be predicted, preparation did not emerge while the mental representation of the item remained latent. Instead, response preparation appeared only after the first response was completed, when the second-tested item became relevant for the upcoming test.

**Figure 5.**
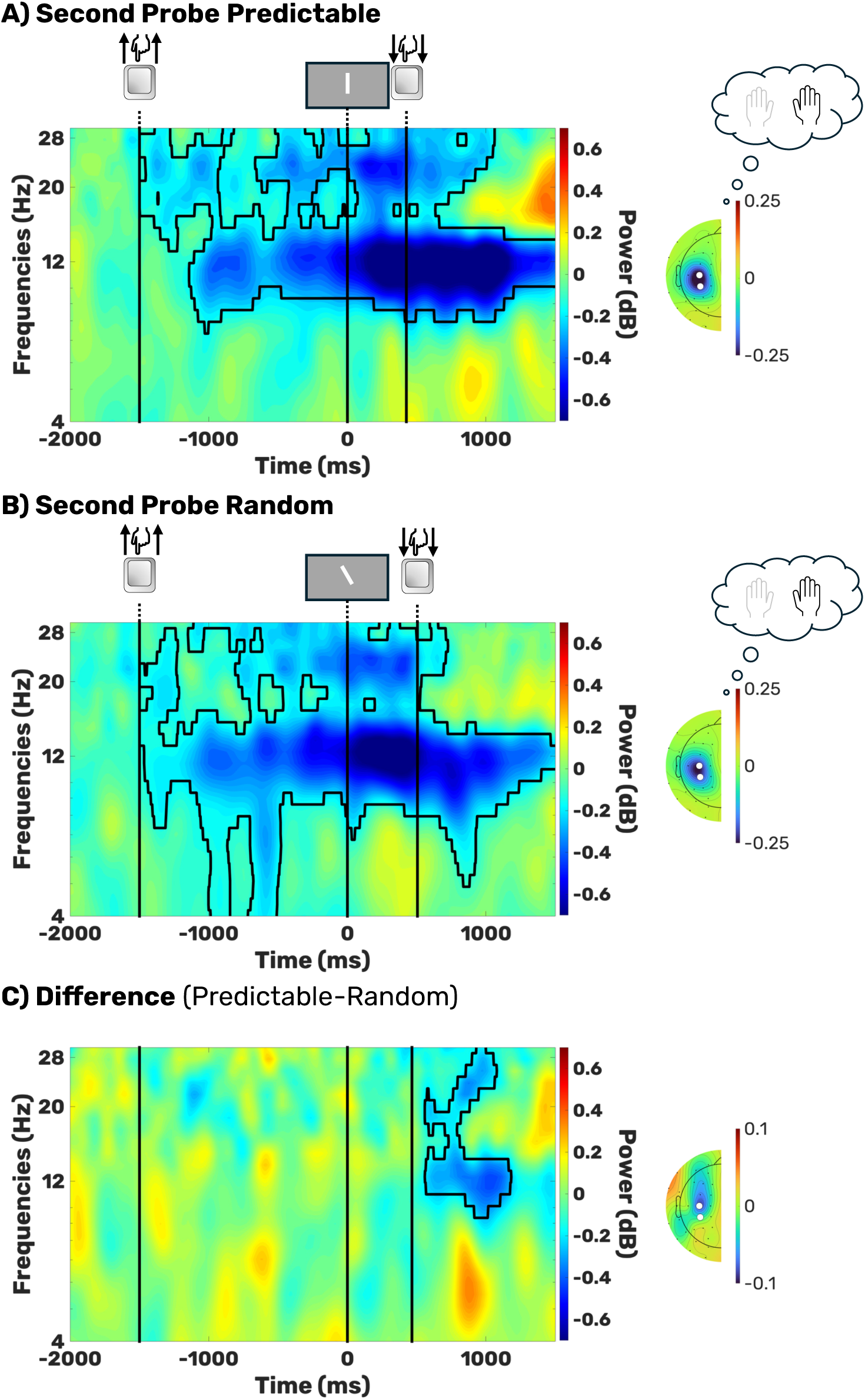
Second response planning following the first response completion. The figure shows lateralization of mu/beta (8–30Hz) EEG power with respect to the tilt-matching hand (channels: C3, C4, CP3, and CP4). Analyses were time-locked to the onset of the second probe and CBPT covered the interval from −1500 to 1500ms. The first black vertical line indicates release of the response button for the first probe, the second black vertical line indicates onset of the second probe, and the third black vertical line indicates the average response onset (button press) for the second probe. In panels A and B, the response-onset line represents the average within-condition response onset, whereas in panel C it represents the average response onset across all trials. Regardless of whether the second probe was predictable (A) or random (B), significant clusters of EEG power lateralization emerged with respect to the prepared response hand. C shows a difference in lateralization between the predictable and random probe conditions following the response. The results show a similar pattern to the first probe window.

Response preparation for the second-tested item showed the same pattern observed for the first-tested item: before probe onset, participants prepared the tilt-matching hand regardless of probe type, whereas differences between predictable and random probes emerged only after the probe appeared. Specifically, following second-probe onset, mu/beta lateralization differed between predictable and random probes (Figure 5C; 560–1180ms, 9.7–30Hz; cluster size = 988, dmax = −0.801, davg = −0.506). As with the first probe, this post-probe difference indicates that when the given probe was random, participants adjusted the initially prepared hand after seeing the random probe’s initial position, selecting the response that provided the more convenient rotation toward the target orientation.

### WM can offload features of visual information into action-related motor codes

Across both encoding and maintenance, we observed preparation of the tilt-matching hand regardless of probe type, even though random probes provided no anticipatory information about the ultimately required response. We therefore tested whether this prepared hand also biased participants’ actual response. Across trials, participants used the tilt-matching hand in 93.39% of predictable first probes (SD=2.863); 62.013% of random first probes (SD = 9.914); 84.207% of predictable second probes (SD=7.261); 62.469% of random second probes (SD= 9.233). As expected, tilt-matching hand use was higher for predictable than random probes (*F*(1,32) = 220.368, *p* < .001; *η2* = 0.807; *BF_incl_* = 1.212x10^13^). It was also more frequent for the first than the second test (*F*(1,32) = 46.983, *p* < .001; *η2* = 0.022; *BF_incl_* = 453.033), with a significant interaction between probe type and test order (*F*(1,32) = 70.163, *p* < .001; *η2* = 0.027; *BF_incl_* = 1.178 x 10^9^). For predictable probes, use of the required tilt-matching hand was higher in the first than in the second test (*t*(32) = 10.702; *p_holm_* < .001; *d* = 1.176; 95% CI [6.851 11.528]).

Critically, using the tilt-matching hand in random-probe trials remained above chance for both the first (*t*(32) = 6.961; *p* < .001; *d* = 1.212; 95% CI [0.755 1.658]), and second random probes (*t*(32) = 7.758; *p* < .001; *d* = 1.350; 95% CI [0.871 1.819]). Thus, the prepared hand biased the subsequent response hand, but did not determine it. During maintenance, mu/beta lateralization did not differ between predictable and random probes, indicating that participants initially prepared the same tilt-matching hand in both random and predictable probes. After probe onset, however, lateralization diverged between probe types for both the first (Figure 4C) and second test (Figure 5C), consistent with participants reevaluating the prepared response once the random probe appeared and switching to the hand that provided the shortest rotation toward the target. Together, the behavioral and EEG results suggest that this preparation functioned as an offloading strategy, with the selected hand carrying information about the item’s tilt direction.

### Predictable action information reduces the need for attentional prioritization on the WM item for the second test

We next examined whether predictability of the upcoming probe reduced the need to allocate selective attention to the tested WM item in both probe windows. Here, the critical information provided by the predictable probe was not which hand to use, but how far the remembered orientation would need to be rotated to match the probe. To index attentional allocation, we calculated posterior alpha lateralization by comparing oscillatory activity over posterior channels contralateral versus ipsilateral to the item’s screen position (see Materials and Methods). For the first-tested item during the first probe window, alpha lateralization did not differ between predictable and random probes. For the second-tested item in the second probe time window, participants reprioritized the item after completion of the first response both when the second probe was predictable (Figure 6A; between -1980- -1168ms and 7.6-13.4Hz; cluster size = 1025, *d_max_* = -0.698; *d_avg_*= -0.474) and when it was random (Figure 6B cluster 1; between -1956- - 1312ms and 7-14.5Hz; cluster size = 1260, *d_max_* = -1.110; *d_avg_*= -0.616). For the second random probe, we observed this visual prioritization reoccurring just before the probe onset (Figure 6B cluster 2; between -412-148ms and 5.5-11.4Hz; cluster size = 867, *d_max_* = -0.838; *d_avg_*= -0.544). Critically, this second attentional reallocation was absent when the second probe was predictable, compared to when it was random (Figure 6C; between -380-148ms and 6.5-11.4Hz; cluster size = 695, *d_max_* = -0.741; *d_avg_*= -0.466). This pattern suggests that probe predictability reduced the demands on the selective attention towards the corresponding item itself for the second-tested item, with knowledge of the precise required rotation partly supporting performance without relying as strongly on the visual WM item.

**Figure 6.**
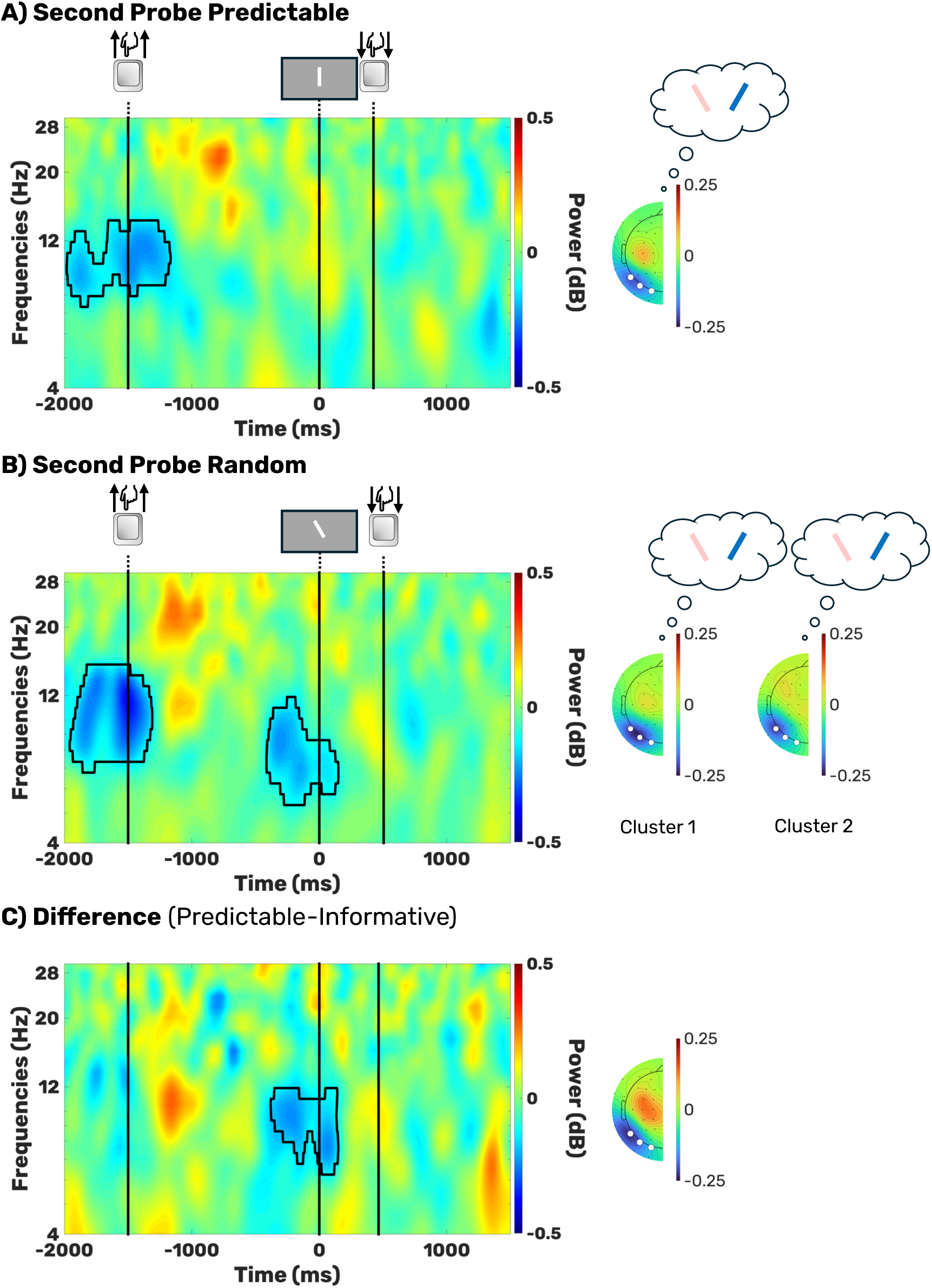
Attentional reprioritization of the latent item for the second test. The figure shows alpha power lateralization (∼8–12Hz) with respect to the target item’s lateral position on the screen (channels: O1, O2, P7, P8, PO7, and PO8). Analyses were time-locked to the onset of the second probe and covered the interval from −1500 to 1500ms. The first black vertical line indicates release of the response button for the first probe, the second black vertical line indicates onset of the second probe, and the third black vertical line indicates the average response onset (button press) for the second probe. In panels A and B, the response-onset line represents the average within-condition response onset, whereas in panel C it represents the average response onset across all trials. (A) When the second probe was predictable, reprioritization of the second test item emerged following completion of the first response. (B) When the second probe was random, reprioritization was also evident around the onset of the second probe. (C) The difference between conditions suggests that when participants could rely on a motor plan based on the predictable probe position, less visuospatial attention was directed toward the target item.

## Discussion

Our study investigated the relationship between action planning and selective attention in visual WM. When the retro-cue prioritized one item for the first test, hand-specific preparation followed the tilt of that item. By contrast, when the predictable probe was assigned to the second test, no response preparation for the latent item was evident after the retro-cue, even though its upcoming action could already be anticipated. This pattern is consistent with previous work showing that latent WM states can preserve information (Rose et al., 2016; Stokes, 2015; Wolff et al., 2017), whereas operations on WM content depend on prioritization of the relevant representation (Trübutschek et al., 2019).

This pattern aligns with Olivers and Roelfsema’s (2020) proposal that prioritization within WM reflects the matching of a memory representation to an upcoming task or action. Within this framework, “action” encompasses both overt motor responses and covert mental operations, thereby determining an item’s relevance to the current task. Our findings support this view by showing that participants did not prepare the response associated with the uncued WM item while prioritizing another item, even when the required response was fully predictable for the uncued one but remained unpredictable for the prioritized.

Our task required participants to learn a mapping between stimulus tilt and response hand, allowing them to anticipate the required response when the probe was predictable. When both memory items shared the same tilt direction, thereby constraining the possible responses to a single hand, we observed hand-specific response preparation already during encoding, reflected in lateralized mu/beta activity (Figure 3). This effect emerged before the retro-cue, when neither item had yet been prioritized, consistent with previous findings. Nasrawi et al. (2023) previously showed a similar effect, where probe-based response mapping was activated along encoding. But notably, in our study, response preparation occurred regardless of whether the probe’s initial orientation was predictable. This suggests that learned stimulus–response associations may automatically activate corresponding motor representations during encoding, generalizing across task contexts independently of the anticipated response requirements.

A similar generalization of learned stimulus–response associations was evident during later stages of the task. Following both the retro-cue and the first probe, we observed preparation of the response hand associated with the tilt of the prioritized item, regardless of whether the anticipated probe required that response. We propose that WM may exploit learned stimulus–response mappings to offload tilt information onto associated motor representations during the maintenance period, thereby reducing WM load (Risko & Gilbert, 2016). This interpretation is consistent with the view that WM maintenance is not restricted to sensory representations but can also recruit other domains, including the motor system (Schneider et al., 2017; Yang et al., 2026). By representing stimulus information through associated motor responses, WM may distribute the maintenance of task-relevant information across sensory and motor systems, potentially reducing reliance on the sensory WM to complete the task.

A complementary form of flexibility emerged from probe predictability. When the second-tested item was paired with a predictable probe, participants could anticipate the precise amount of rotation required before that item became task-relevant. In this condition, posterior alpha lateralization for the latent item was reduced (Figure 6), suggesting that the WM item didn’t need to be prioritized by selective attention as long as the random condition (Foster et al., 2016; Ikkai et al., 2016; Woodman et al., 2022); This indicates that knowing the upcoming precise action required by the anticipated probe position reduced reliance on attentional reprioritization when the second-tested item was to be reported. The behavioral results supported this interpretation: the performance benefit of predictable probes emerged specifically when they were presented second, after the corresponding item had remained unprioritized during the initial maintenance period. Together, these findings suggest that precise prospective information about the required motor response can support latent WM representations by reducing demands on subsequent attentional selection.

In summary, our findings suggest that response preparation is coupled with attentional prioritization in WM. Even when uncued (unprioritized) information in WM requires a predictable action, this response is not prepared unless the WM item becomes prioritized again. This indicates that prioritization within WM remains critical for guiding memory-based actions, rather than being bypassed by the availability of anticipated motor codes. At the same time, our findings highlight the flexibility of WM in recruiting motor representations to support the maintenance of visual information. Having a precise action plan can reduce the need for bringin the WM item back into focus of attention. Furthermore, through learned stimulus–response associations, WM may partially offload information onto the motor system, thereby reducing WM demands when possible. Together, these findings suggest that action planning is not merely a downstream consequence of memory retrieval but may also contribute to how WM organizes and maintains task-relevant information.

## Data Availability

All data and materials will be shared publicly on OpenNeuro upon publication. Only anonymized data will be shared.

## Code Availability

All scripts related to data analysis and experimental setup will be shared publicly on GitHub upon publication.

## Preregistration

The experimental design, data collection plan, hypotheses, and analysis plan for this study have been preregistered on the Open Science Framework prior to data collection (https://doi.org/10.17605/OSF.IO/J8XUB).

## Author Contributions

All authors contributed to the conceptualization of this study and the design of the experiment. Ş.Ö. collected the data and conducted the data analysis. All authors discussed the interpretation of the results and wrote the manuscript.

## Materials and Methods

The experimental design, target sample size, data collection plan, hypotheses, and analysis plan for this study have been preregistered on the OSF before data collection (LINK).

### Participants

Thirty-four participants voluntarily took part in the study. One participant was excluded from further analysis due to their performance being 2 standard deviations (SD) away from all participants’ mean (for preregistered exclusion criteria, see OSF). One participant’s EEG data was excluded due to excessive noise; however, behavioural data were included. Final analyses were conducted on 32 EEG datasets and 33 behavioural datasets.

Our primary hypothesis concerns memory-guided response planning. To test it, we quantified mu/beta power lateralization relative to the designated response hand. In a previous study, we observed robust mu/beta power lateralization with a mean effect size of Cohen’s d = 0.726 across the significant cluster identified by a cluster-based permutation test (CBPT; Özdemir et al., 2025). However, the present study introduces a novel approach to investigating the temporal dynamics of action planning. Because the effect sizes for these, such as H1, which focuses on the encoding window and predicts action planning based on stimulus characteristics, could not be estimated reliably, we adopted a more conservative approach to determining the sample size. An a priori power analysis conducted in G*Power (Faul et al., 2007) for a one-tailed paired-samples t-test, assuming *d* = 0.726, α = .05, and power = .95, indicated a required sample size of 22 participants. To account for the possibility of a smaller and less reliable effect in the present study, we also considered a medium effect size of *d* = 0.50, which yielded a required sample size of 45 participants. Given the available infrastructure, time constraints, and personnel resources, we based our final estimate on the midpoint between the previously observed and medium effect sizes *d* = 0.61). This resulted in a target sample size of 32 participants. This sample size is also comparable to those used in previous studies investigating working-memory-related response preparation.

### Ethics Statement

This study was conducted under the ethics approval of Leibniz Research Centre for Working Environment and Human Factors and was in accordance with the Declaration of Helsinki.

### Apparatus, stimuli, and procedure

The experiment was conducted using a ViSaGe MKII Stimulus Generator. Stimuli were presented on a 22-inch CRT monitor at a resolution of 1024 × 768 pixels. The refresh rate of the screen was 100Hz. Participants completed the experiment in a dimly lit, electrically shielded room. The paradigm was programmed in Free Pascal using the Lazarus integrated development environment. Responses were collected via keyboard button presses.

Participants were asked to complete a handedness questionnaire (Oldfield, 1971) to assess their dominant hand. All participants were classified as right-handed. Color vision was evaluated via the Ishihara test; none of the participants showed signs of color-blindness.

Stimulus orientations were sampled from a uniform distribution ranging from 0° to 180°, excluding orientations between 85° and 95° to avoid near-vertical stimuli. Orientations were then pseudorandomized across trials. Within each trial, the orientations of the two items differed by at least 10°. The design included equal numbers of trials in which both items were tilted to the left, that is, counterclockwise from vertical, and both items were tilted to the right, that is, clockwise from vertical. Tilt direction was further counterbalanced across items presented on the left and right sides of the display. Trials with both items matching and nonmatching tilt directions were equally represented. The experiment consisted of 1,200 trials in total.

In each trial, participants were first presented with two tilted lines (3.5° × 0.35°) for 250ms and instructed to remember their orientations. One line was blue (CIE 1931: 0.193, 0.221, 40), and the other was orange (CIE 1931: 0.500, 0.432, 40). A white fixation point (diameter: 0.33°) was then presented for 800ms, followed by a retro-cue for 250ms. The retro-cue matched the color of one of the memorized items, indicating that this item would be tested first. After a 1,500-ms fixation interval, the first probe was presented to test the cued item.

Participants were required to initiate their response within 2,000ms. They used the “A” key to rotate the probe counterclockwise and the “L” key to rotate it clockwise. While a key was held down, the probe rotated continuously at a rate of 0.6° per frame. When the participant released the key, the final orientation of the probe was recorded as the response. Following key release, a fixation point was presented for 1,500ms. The second probe was then presented to test the non-cued, latent item, using the same response procedure. Inter-trial interval varied between 800 and 1200ms.

In all trials, either the first or the second probe initially appeared in the vertical position. This was manipulated blockwise. Before each block, participants were informed which probe (the first or the second) would be presented vertically, allowing them to prepare the corresponding response in advance. We refer to this probe as the predictable probe.

### EEG Preprocessing

EEG preprocessing was performed in MATLAB (R2023b; MathWorks) using the ERPLAB (Lopez-Calderon & Luck, 2014) and EEGLAB (Delorme & Makeig, 2004) toolboxes. First, the data were high-pass filtered at 0.1Hz and low-pass filtered at 40Hz using an IIR Butterworth filter implemented with ERPLAB’s ‘pop_basicfilter’ function [default order: 3 × (sampling rate / low cutoff)]. Channels containing excessive artifacts were then automatically identified and rejected using EEGLAB’s ‘pop_rejchan’ function with a kurtosis threshold of 15. After the automated rejection of channels, the data were re-referenced to the average of all remaining channels.

To prepare the data for independent component analysis (ICA), the processing steps below were performed after re-referencing. After the independent-component weights had been computed, the pipeline returned to the re-referenced dataset, and the ICA weights were applied to these data without retaining the intermediate preprocessing steps. The steps in between took place only to prepare the dataset for ICA.

ICA-specific steps: After re-referencing, the data were downsampled to 200Hz and high-pass filtered with a 1Hz threshold using a Hamming-windowed sinc FIR filter (pop_eegfiltnew; filter order, 661; transition bandwidth, 1Hz; cutoff frequency at −6 dB, 0.5Hz). Then, the data were epoched starting 600ms before stimulus onset and ending 8300ms after. Baseline correction was performed relative to the 200ms time window prior to stimulus onset. Noisy trials were rejected using EEGLAB’s pop_autorej function (threshold, 500 µV; maximum percentage of rejected trials per iteration, 5). Then, ICA was conducted following trial rejection. Independent components (ICs) were then identified using the ICLabel classifier of EEGLAB, classifying components into categories such as the brain, muscle, eye movements, heart activity, channel noise, line noise, and “other” (Pion-Tonachini et al., 2019). ICs with a probability of being eye movement >30% and with a probability of being brain signal <10% were marked for rejection.

ICs identified as artifacts by the abovementioned criteria were removed, and the remaining IC information was attached to the rereferenced data, bypassing the intermediate ICA-preparation stages. The continuous data were then epoched from −600 to 8300ms relative to stimulus onset, defined as time zero. Baseline correction was performed using the 200-ms prestimulus interval. Trial rejection was conducted with EEGLAB’s pop_autorej function (threshold, 500 µV; maximum percentage of rejected trials per iteration, 5). This trial rejection was the final rejection for the data (abovementioned rejection for ICA wasn’t carried out after IC rejection). Any remaining noisy trials were rejected using the same automated procedure applied during ICA preparation. Finally, missing channels were interpolated using EEGLAB’s ‘pop_interp’ function with spherical spline interpolation, and the data were downsampled to 250Hz.

Our preprocessing pipeline resulted in rejecting 27.16 ICs on average (SD = 5.17). Excluding the blink artifacts (which are large deflections), the rejected ICs explained on average 9.68% of the variance across all participants (SD = 8.05), including components related to muscle activity, heartbeats, line noise, channel noise, and other artifacts. As a result of artifact rejection, we had an average of 986.58 remaining trials (SD = 6.36). Remaining trial counts per condition were as follows: Predictable Probe First (mean = 499.38; SD = 31.65), Predictable Probe Second (mean = 487.19; SD = 29.68).

### EEG Analysis

We first calculated the time–frequency (TF) decomposition. For each trial and channel, the oscillatory power for each frequency was calculated from 4 to 30Hz on a logarithmic scale in 26 steps. For each frequency in the given spectrum, a complex Morlet wavelet was created by tapering a sinusoid (ei2ft) of the given frequency with a Gaussian [(e− t2/2s2; s is the width of the Gaussian; s = /(2f ) corresponds to the number of cycles of the wavelet)]. Our zero-padded data and the Morlet wavelets were fast Fourier transformed (FFT). Convolution in TF decomposition was done by multiplying the signal’s FFT with the FFT of a complex Morlet wavelet at each frequency by taking the dot product. Then an inverse FFT was applied to get the time-domain result. We applied a baseline normalization by averaging the −400ms to −100ms time interval in all trials and divided each trial by this averaged value and converted the measurement to decibel (dB).

Our primary hypotheses concerned motor planning and hand preparation based on WM representations. To examine these processes, we measured lateralization of mu/beta-band power (∼8–30Hz) over centroparietal EEG electrodes (C3, C4, CP3, and CP4; Boettcher et al., 2021; Özdemir et al., 2025; Rösner et al., 2022; Schneider et al., 2017) relative to the prepared response hand. Lateralization was calculated by subtracting ipsilateral from contralateral power. Mu/beta lateralization indexes response preparation by reflecting preparatory activity associated with the corresponding hand.

Exploratory analyses examined the allocation of internal visual attention. We calculated lateralization of posterior alpha-band power (∼8–12Hz; Foster et al., 2016; Ikkai et al., 2016; Woodman et al., 2022) relative to the screen position of the relevant item using electrodes Oz, O1, O2, Pz, PO3, and PO4. As for mu/beta power, alpha lateralization was calculated by subtracting ipsilateral from contralateral power.

To test whether these lateralization effects emerged within the specified time windows or differed between conditions, we used cluster-based permutation tests (CBPTs). At each time– frequency point, contralateral-minus-ipsilateral power was compared with zero using a one-sample t-test. For condition comparisons, the difference between the two lateralized measures was compared with zero using a one-sample t-test. Points meeting an uncorrected threshold of p < .05 were grouped into candidate clusters. We then performed 1,000 sign-flip permutations by randomly multiplying each participant’s difference scores by either −1 or 1. For each permutation, we recorded the size of the largest significant cluster to generate a null distribution. Observed clusters exceeding the 95th percentile of this distribution were considered statistically significant. For each significant cluster, Cohen’s d was calculated at every time–frequency point, and the maximum and mean effect sizes across the cluster were reported.

### Behavioral Analyses

Behavioral error was calculated for each trial as the absolute angular difference between the participant’s response and the target orientation. Performance in each condition was quantified for each participant by averaging these absolute angular differences.

To determine whether the observed effects on behavioral performance were influenced by any response bias that can be caused by the anticipatory differences between the two probe types, we repeated the analysis using only random-probe trials in which participants selected the shortest rotation direction to the target orientation (Figure 2A). This analysis tested whether the advantage of predictable probes was driven by participants being guided toward the shortest rotation direction by the vertical appearance of the predictable probe. As described above, absolute angular errors were averaged for each condition and participant.

Furthermore, to quantify the bias toward using the tilt-matching hand, we calculated, for each participant, the percentage of trials in which the tilt-matching hand was used in the random probe and predictable probe conditions.

Lastly, we calculated the response onset times for each response and averaged these within each condition per dataset. The onset of the response was defined by the first button press after memory probe presentation.

For all four analyses, condition differences were assessed using a 2 × 2 within-subjects ANOVA, with test order (1st probe vs. 2nd probe) and whether the starting position of the probe is predictable (random probe vs. predictable probe) as within-subject factors. For main effects, eta-squared (η²), for post-hoc analyses, Cohen’s d was reported as the effect size. For all behavioral analyses, both frequentist and Bayesian statistics were reported. In Bayesian repeated-measures ANOVAs, the BF₁₀ value represented the comparison between the tested model, which included the relevant main effect, and the null model. For interaction analyses, the null model included the main effects (Bergh et al., 2020). All Bayesian analyses used JASP’s default priors (Bergh et al., 2020; Love et al., 2019). All p-values from multiple comparisons following the ANOVA were adjusted using the Holm method.

